# Crystal structure of *Bifidobacterium bifidum* glycoside hydrolase family 110 α-galactosidase specific for blood group B antigen

**DOI:** 10.1101/2024.03.03.583176

**Authors:** Toma Kashima, Megumi Akama, Takura Wakinaka, Takatoshi Arakawa, Hisashi Ashida, Shinya Fushinobu

## Abstract

To overcome incompatibility issues and increase the possibility of blood transfusion, technologies that enable efficient conversion of A- and B-type red blood cells to the universal donor O-type is desirable. Although several blood type-converting enzymes have been identified, detailed understanding about their molecular functions is limited. α-Galactosidase from *Bifidobacterium bifidum* JCM 1254 (AgaBb), belonging to glycoside hydrolase (GH) 110 subfamily A, specifically acts on blood group B antigen. Here we present the crystal structure of AgaBb, including the catalytic GH110 domain and part of the C-terminal uncharacterized regions. Based on this structure, we deduced a possible binding mechanism of blood group B antigen to the active site. Site-directed mutagenesis confirmed that R270 and E380 recognize the fucose moiety in the B antigen. Thermal shift assay revealed that the C-terminal uncharacterized region significantly contributes to protein stability. This region is shared only among GH110 enzymes from *B. bifidum* and some *Ruminococcus* species. The elucidation of the molecular basis for the specific recognition of blood group B antigen is expected to lead to the practical application of blood group conversion enzymes in the future.

## INTRODUCTION

The A and B antigens of the ABO(H) blood group system are oligosaccharides present on the surface of human red blood cells that play a key role in determining compatibility of blood for transfusions.^1–3)^ Blood group antigen epitopes are also present in mucin *O*-glycans in the intestinal mucosa.^4)^ The blood group O antigen (H antigen) is composed of the trisaccharide Fuc-α1,2-Gal-β1,3-GlcNAc. The A and B antigens differ from H antigen by the addition of an extra sugar moiety, this being α1,3-linked GalNAc for A-type and Gal for B-type, to the β1,3-linked Gal residue (Figure 1A).^5)^ Because type O blood can theoretically be injected into patients with any of the ABO blood types, it is often considered a universal donor blood type. Attempts are being made to convert A and B antigens to the H antigen by hydrolyzing the GalNAc-α1,3- and Gal-α1,3-glycosidic linkages. The use of blood group converting enzymes, as exemplified by the recently discovered system using a combination of deacetylase and glycoside hydrolase (GH) family 36 α-galactosaminidase, is considered the most promising method for removing the desired sugars.^6,7)^ In 2007, Liu *et al*. discovered two novel GH families that specifically convert type A and B antigens to type H antigen.^8)^ This discovery led to the establishment of GH109 α-*N*-acetylgalactosaminidase and GH110 α-galactosidase families in the carbohydrate-active enzyme (CAZy) database.^9)^ GH110 utilizes an anomer-inverting mechanism and is classified into three major subfamilies (Figure 1B). Members of the GH110 subfamily A specifically act on branched blood group B antigen (Gal-α1,3-(Fuc-α1,2-)Gal), whereas those of subfamily B do not.^10)^ The third unclassified subfamily contains an α-galactosidase from *Pseudoalteromonas distincta* U2A (PdGH110B).^11)^ PdGH110B shows activity toward α1,3-linked galactobiose and was postulated to act on Gal-α1,3-Gal2S released from λ-carrageenan. However, PdGH110B was not active toward the branched blood group B antigen. Structural analysis of PdGH110B revealed the catalytic acid residue as D344, whereas the catalytic base residue could not be identified as D321 or D345.^11)^ Currently, the only GH110 enzyme whose three-dimensional structure has been reported is PdGH110B; the three-dimensional structures of enzymes belonging to subfamilies A and B remain unknown.

**Figure 1.**
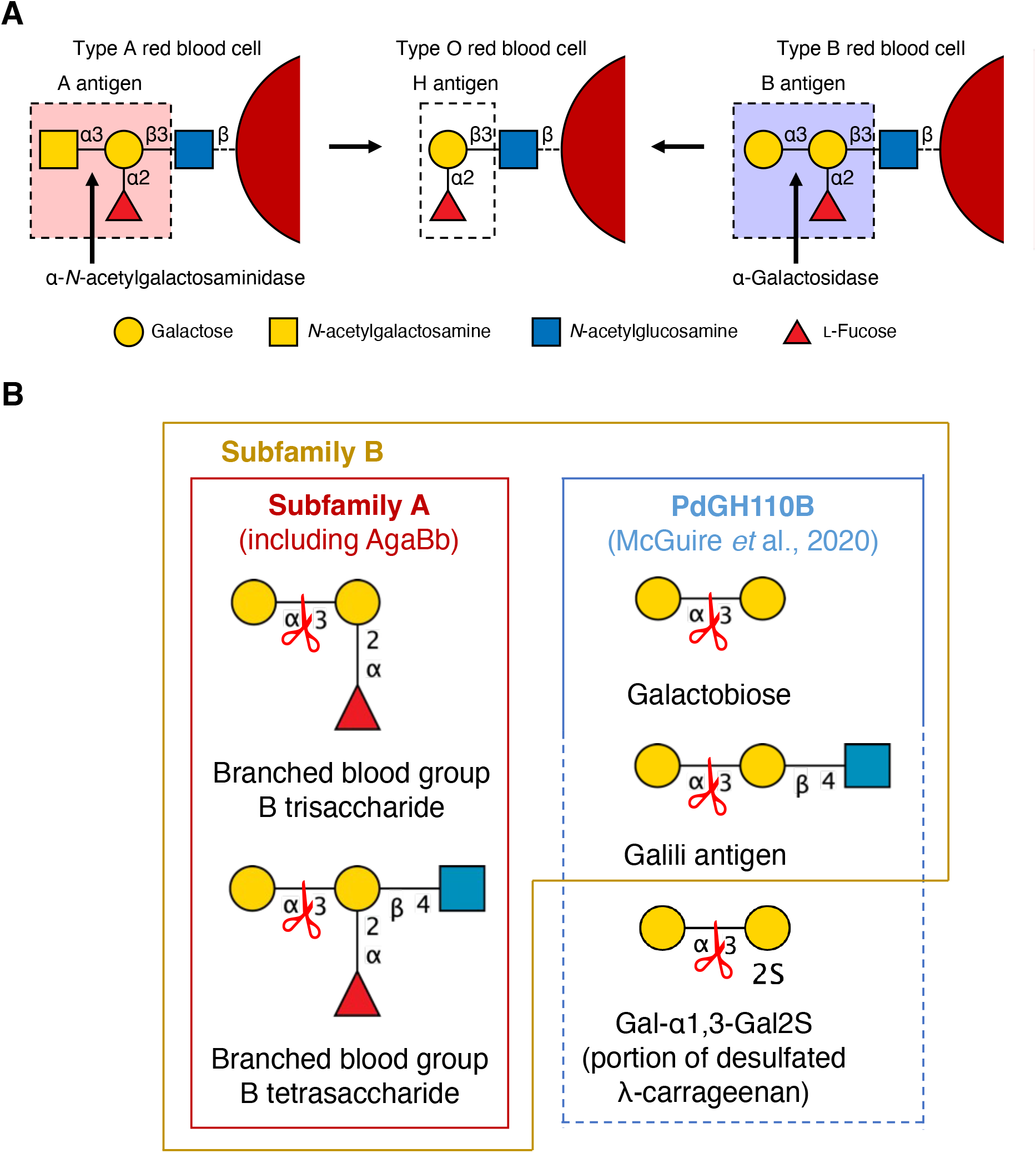
Enzymatic blood group conversion and substrate specificities of GH110. (A) Glycan structure of blood group epitopes and the enzymes implicated in the conversion of blood group A and B epitopes into universal O blood group. (Prepared based on Rahfeld *et al*., 2019).^39)^ (B) Differences in substrate specificity between GH110 subfamilies. Cleavage sites of the enzymes are indicated with red scissors. Substrates that PdGH110B is expected to cleave, but remain to be experimentally verified are indicated with a dashed line box.

Bifidobacteria are common inhabitants of the animal gastrointestinal tract. Infant gut-associated bifidobacteria, including *Bifidobacterium bifidum*, exert various beneficial effects on human health.^12)^ *B. bifidum* assimilates mucin *O*-glycans and possesses many cell surface-anchored GHs that act on *O*-glycans.^13)^ Wakinaka *et al*. discovered an α-galactosidase from *B. bifidum* JCM 1254 (AgaBb), which belongs to GH110 subfamily A and specifically acts on blood group B antigen.^14)^ AgaBb, together with other membrane-bound GHs, has also been implicated in the degradation of the blood group B antigen present at the non-reducing end of core 2 *O*-glycan epitope of the intestinal mucosa.^13)^ In this study, we determined the crystal structure of AgaBb, including the GH110 catalytic domain and part of the C-terminal uncharacterized domain. We also performed mutational analysis of the active site residues and thermal shift assays on constructs of various lengths to identify the substrate-recognition mode and domain responsible for the thermostability of AgaBb. While preparing this paper, the crystal structure of α-galactosidase active on blood group B antigen was released, so we also compared the AgaBb structure with it in the Discussion section.

## MATERIALS AND METHODS

### Materials and chemicals

Unless otherwise indicated, all the reagents were purchased from Fujifilm Wako Pure Chemicals (Osaka, Japan).

### Production and purification of recombinant protein

Constructs used for heterologous protein expression were derived from a pET23b construct with full-length AgaBb described previously.^14)^ A schematic representation of each construct is shown in Figure S1. The characteristics and design of these constructs are summarized in Table S1. For the expression of T7-tagged AgaBb700, a plasmid previously prepared to produce CBM51-deleted AgaBb was used. The other constructs were prepared using a modified sequence deletion method with complementary primers.^15)^ PCR was performed using KOD One PCR Master Mix (Toyobo, Osaka, Japan). The PCR product was treated with DpnI (TaKaRa Bio) at 37 °C for 1 h and transformed into *Escherichia coli* JM109. The primers used for sequence editing along with the complementary primers are listed in Table S2. *E. coli* Rosetta2 (DE3) (Merck Millipore, Darmstadt, Germany) was transformed with each expression vector and cultured in lysogeny broth medium containing 100 μg/mL ampicillin and 17 μg/mL chloramphenicol at 37 °C until the OD_600_ (optical density at 600 nm) reached 0.4–0.6. Gene expression was induced by adding isopropyl-β-D-thiogalactopyranoside at a final concentration of 0.1 mM, and the culture was continued for another 20 h at 25 °C. Bacterial cells were harvested by centrifugation and suspended in a lysis buffer (50 mM Tris-HCl (pH 7.0) and 200 mM NaCl). The cell suspension was sonicated using a 250D sonicator (Branson Ultrasonics Corporation, Danbury, CT, USA). The lysate was centrifuged, and the supernatant was loaded on a column filled with cOmplete His-tag Purification Resin (Roche Diagnostics GmbH, Mannheim, Germany) and pre-equilibrated with lysis buffer. After loading the crude sample, the column was washed with lysis buffer supplemented with 5 mM imidazole and the protein was eluted with lysis buffer supplemented with 500 mM imidazole. The eluate was then concentrated and desalted using Amicon Ultracel-10 kDa Centrifugal Filters (Merck Millipore). The sample was loaded onto a Mono Q 10/100 GL column (Cytiva, Marlborough, MA, USA), pre-equilibrated with 25 mM Tris-HCl (pH 7.0), and subjected to a linear gradient of NaCl concentrations up to 0.5 M. The target protein was eluted with approximately 200 mM NaCl. The sample fractions were concentrated using Amicon Ultracel-10 kDa Centrifugal Filters and loaded onto a HiLoad 16/60 Superdex 200 pg column (Cytiva) pre-equilibrated with 25 mM Tris-HCl (pH 7.0) and 200 mM NaCl. Approximately 20–50 mg of the target protein (>99% purity) per liter of culture was obtained after sequential column purification steps. Fractions corresponding to the target proteins were concentrated, and the buffer was exchanged with 10 mM Tris-HCl (pH 7.0) and 200 mM NaCl using Amicon Ultracel-10 kDa centrifugal filters. The purified protein (>40 mg/mL concentration) was stored at 4 °C until further use. Purity of the protein sample was verified by sodium dodecyl sulfate polyacrylamide gel electrophoresis (SDS-PAGE) at the purification and preservation steps, and analyzed using GelAnalyzer (version 19.1). Protein concentration was determined by bicinchoninic acid assay using bovine serum albumin (TaKaRa Bio) as the reference standard. For the crystallized sample, which required a higher concentration, the protein concentration was determined by measuring the absorbance at 280 nm using a Nanodrop 2000 spectrophotometer (Thermo Fisher Scientific, Waltham, MA, USA). The molar extinction coefficient used in this study was calculated for each construct from the amino acid sequence using the ProtParam tool in Expasy (https://web.expasy.org/protparam/).

### Protein crystallography and substrate modeling

Crystallization screening was performed for every construct (Figure S1) using the sitting-drop vapor-diffusion method at 20 °C by mixing equal volumes of the protein solution with the reservoir solution in the JCSG Core I–IV Suite Kit (NeXtal Biotechnologies, Holland, OH, USA). In addition to the conventional optimization of crystallization conditions, microseed matrix screening was performed using an in-house kit inspired by the PEGs Suite Kit (NeXtal Biotechnologies) with a protein: reservoir: seed ratio of 6:4:2.^16)^ Among the many crystallization hits obtained, the three conditions under which diffraction-quality crystals were obtained are listed in Table S3. Co-crystals were prepared using a protein solution containing 20 mM D-galactose. Crystals used for the single-wavelength anomalous diffraction (SAD) method were prepared by soaking the crystals of T7-tag_AgaBb700 in a reservoir containing 5 mM K_2_PtCl_4_ for 28 h. The crystals were cryoprotected in a reservoir solution supplemented with a cryoprotectant and flash-cooled by immersing in liquid nitrogen.

Diffraction data were collected from the beamlines at the Photon Factory of the High-Energy Accelerator Research Organization (KEK, Ibaraki, Japan). The diffraction datasets were processed using the XDS^17)^ and Aimless.^18)^ The phase resolution was determined using the CRANK2 pipeline,^19)^ and molecular replacement was performed using Phaser.^20)^ Automated model building was performed using Buccaneer ^21)^. Refinement was performed using PHENIX.^22,23)^ Manual model rebuilding was performed using Coot.^24)^ Molecular graphics were prepared using PyMOL (Schrödinger LLC, New York, NY, USA). A model of blood group B trisaccharide was designed using the Glycam carbohydrate builder tool (https://glycam.org/cb/) and manually placed in the active site of AgaBb by superimposing the Gal-α1,3-Gal group onto the galactobiose molecule in the complex structure of PdGH110B (PDB ID: 7JWF). The model was energy-minimized using the PyMOL sculpting wizard. All the pyranose rings in the models were chair conformations.

### Thermal shift assay

Thermal stability was verified using the StepOne Real-Time PCR system (Thermo Fisher Scientific Japan, Tokyo, Japan) and accompanying system control software. SYPROorange dye (1×; Thermo Fisher Scientific) was added to 10, 15, and 20 mg/mL protein solutions. A temperature gradient of 25–99°C was applied and the fluorescence was monitored at 605 nm. The temperature at which the derivative of the fluorescence (in arbitrary units) was the highest was designated as the melting temperature (*T*_m_ value) of the target protein. HEPES-NaOH (0.1 M; pH 7.0) was used as the sample diluent as the pH of Tris-HCl changes with thermal fluctuations. The *T*_m_ data were analyzed by two-way ANOVA using Prism 8 (GraphPad Software, San Diego, CA, USA).

### Site-directed mutagenesis

Site-directed mutagenesis was performed on the AgaBb844 construct using the complementary primer sets listed in Table S2. PCR, DpnI digestion, transformation, protein production, and purification were performed as described in the previous section.

### Enzyme assay

Enzymatic assays were performed as described previously.^14)^ For thin-layer chromatography (TLC), blood group B trisaccharide (1 mM) was incubated with the purified protein (1 μg/mL) in 50 mM Na-acetate (pH 6.0) overnight at 37 °C. 1-Butanol: acetic acid: water (2:1:1, v/v) was used as the developing solvent and orcinol sulfate as the dyeing reagent.^25)^ For galactose dehydrogenase-coupled assay,^26,27)^ blood group B trisaccharide (2 mM; Biosynth Carbosynth, Compton, UK) was incubated with 1 μg/mL of the protein in 50 mM sodium acetate (pH 6.0), and the reaction was stopped by heating at 95 °C for 5 min. The reaction mixture was diluted 10-fold by adding 100 mM Tris-HCl (pH 8.6), 3 mM NAD^+^, 0.4 mg/mL BSA, 2 mM EDTA, and 1 U of galactose dehydrogenase (GalDH, Megazyme, Wicklow, Ireland). The absorbance was monitored at 340 nm until equilibrium was reached. The concentration of NADH produced, which equals the concentration of galactose before the GalDH treatment, was calculated based on the molar extinction coefficient of 6.22 × 10^3^ M^-1^cm^-1^

### Multiple alignment and phylogenetic tree

Multiple sequence alignments were performed using M-Coffee.^28)^ Multiple alignments of the characterized GH110 enzymes were performed using ESPript 3.0.^29)^ Homolog sequences were retrieved using a BLAST search against the reference protein database (RefSeq proteins) in NCBI with default settings for sequence filtering or selection. The domains of the retrieved protein sequences were annotated by integrating the HMMER and DIAMOND tools on the dbCAN3 meta server.^30)^ A phylogenetic tree was constructed based on the maximum-likelihood algorithm with a bootstrap value of 1000 using MEGA^31)^ and displayed using iTOL v6.^32)^

## RESULTS

### C-terminal deletants

AgaBb is a 1289 amino-acid polypeptide containing an N-terminal signal peptide (residues 1–23), a GH110 domain (residues 24–673), an uncharacterized region (residues 674–942), a carbohydrate-binding module (CBM) family 51 domain (CBM51, residues 943–1095), a bacterial immunoglobulin-like group 2 domain (Big2, residues 1103–1184), and a C-terminal transmembrane region (residues 1256–1283) (Figure 2A). The full-length AgaBb construct constructed previously lacked the N-terminal signal peptide and C-terminal transmembrane region, and contained an N-terminal T7-tag and C-terminal 6×His-tag instead (Figure S1).^14)^ In the present study, we generated a construct that lacked the N-terminal T7-tag of the full-length construct (residues 23–1191, AgaBb1191), and five other constructs with stepwise deletion from the C-terminus to the last 637^th^ amino acid in the GH110 domain (AgaBb1099, AgaBb945, AgaBb844, AgaBb730, and AgaBb673). The activity of the purified proteins toward blood group B trisaccharide was confirmed for all six deletant constructs (Figure S2). The molecular weights of the six constructs and one additional construct (AgaBb1180, Figure S1) were examined by size-exclusion chromatography (Figure S3). As in the case of full-length AgaBb,^14)^ the two longer constructs (AgaBb1180 and AgaBb1191) were eluted at positions corresponding to a dimer. In contrast, the five shorter constructs (AgaBb1099, AgaBb945, AgaBb844, AgaBb730, and AgaBb673) were measured as monomers in solution, suggesting that the Big2 domain is involved in dimerization.

**Figure 2.**
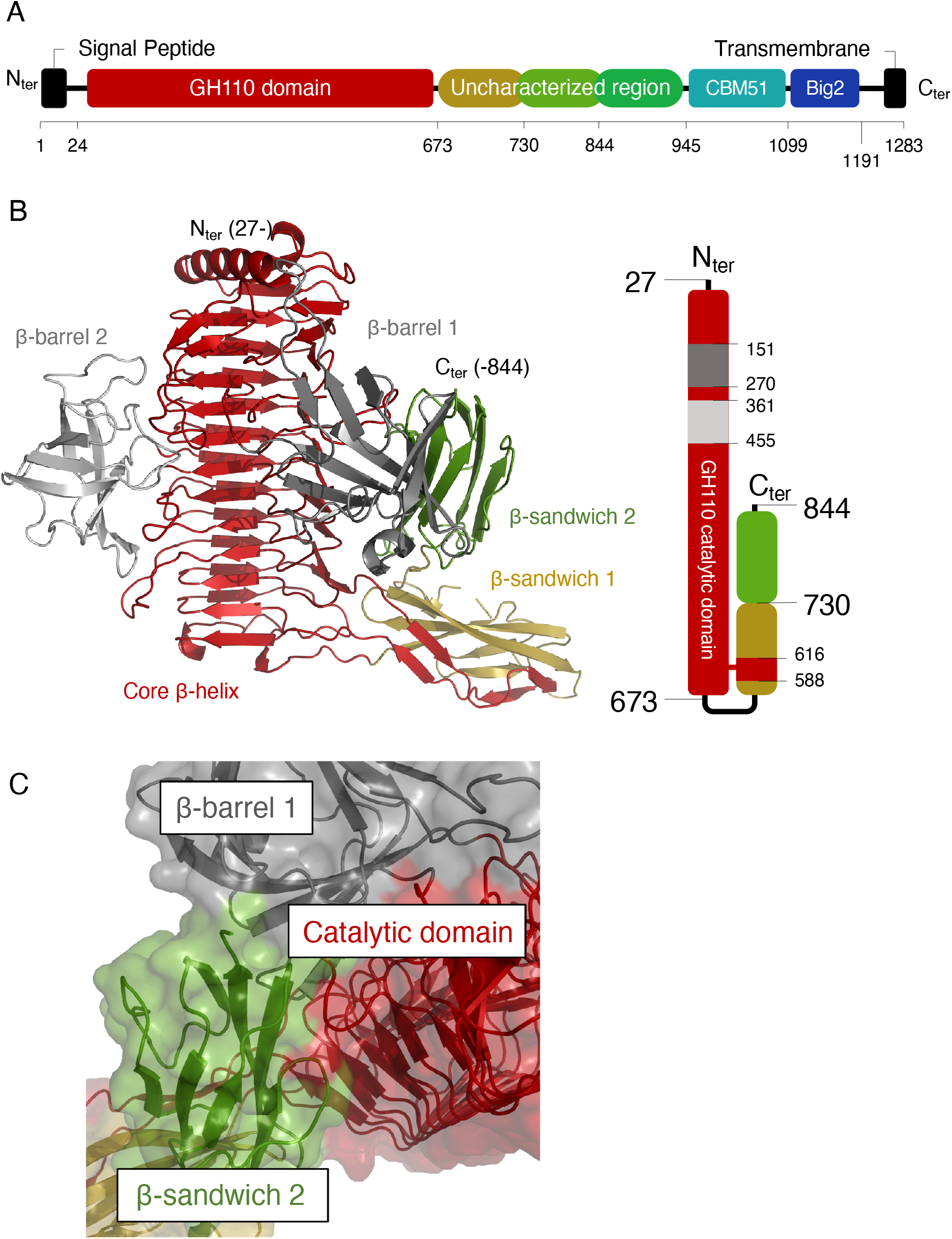
Domain organization of AgaBb and crystal structure of AgaBb844. (A) Domain organization of AgaBb. GH110, glycoside hydrolase family 110. AgaBb, α-galactosidase from *Bifidobacterium bifidum*. (B) The overall structure (left) and domain architecture (right) of AgaBb344. The monomer is colored to show the core β-helix (red), β-barrel 1 (dark gray), β-barrel 2 (light gray), β-sandwich 1 (gold), and β-sandwich 2 (green). (C) Surface representation of the β-sandwich 2 domain showing extensive interactions with the core β-helix and β-barrel 1 domain.

### Crystal structures

An AgaBb construct (residues 24–700) with an N-terminal T7-tag (T7-tag_AgaBb700) was previously used as a CBM51-deleted AgaBb (Figure S1).^14)^ First, we crystallized T7-tag_AgaBb700 and succeeded in initial phase determination using the SAD method at 3.15 Å resolution using a crystal soaked in 5 mM K_2_PtCl_4_ (Table S4). The native protein structure of the T7-tag_AgaBb700 was determined at 1.96 Å resolution. However, the reproducibility of T7-tag_AgaBb700 crystallization was very low. Examination of the crystal structure revealed that the C-terminal region (residues 618–700) was interlaced with the neighboring molecule in the crystal packing, and the preceding region (residues 587–617) was disordered (Figure S4A). The dimeric structure in the crystal packing was not consistent with the monomeric structure measured by size-exclusion chromatography, suggesting that intermolecular interlacement was an artifact that hindered the reproducibility of crystallization. Therefore, we created a construct with further C-terminus deletion (T7-tag_AgaBb673, Figure S1) and crystallized it with 20 mM galactose. The structure of T7-tag_AgaBb673 showed no intermolecular entanglement in crystal packing, and a plausible electron density for galactose was observed at the active site (Table S4 and Figure S4B). However, the X-ray diffraction resolution of the T7-tag_AgaBb673 crystal was very poor (maximum at 3.50 Å) and did not improve even with vigorous crystallization screening. After the construction of C-terminal deletants without the N-terminal T7-tag (Figure S1) and extensive crystallization screening, we succeeded in crystallizing AgaBb844 (residues 23–844), which included a part of the C-terminal uncharacterized region, with good reproducibility. The crystal structure of AgaBb844 was determined at 2.02 Å resolution (Table S4). The catalytic domain structure of AgaBb844 was similar to those of T7-tag_AgaBb700 and T7-tag_AgaBb673 because the average root mean square deviation (RMSD) values for 490 Cα atoms were <0.43 Å. Unless otherwise noted, we mainly describe the AgaBb844 structure because it had the longest structure.

The asymmetric unit of AgaBb844 contained three protein molecules. Protein interfaces, surfaces, and assemblies (PISA) server analysis^33)^ confirmed that the monomeric structure corresponded to a possible biological assembly, as implied by size-exclusion chromatography. Because the three chains in the asymmetric unit have virtually identical structures (Cα RMSD values between all chain pairs are < 0.27 Å), we have mainly described chain A. Almost all amino acids were modeled from residues 27–844, except for the disordered residues 732–739, including the GH110 catalytic domain and a part of the uncharacterized region (Figure 2B). No electron-density map was observed for the T7-tag. The GH110 domain consists of a core right-handed parallel β-helix of 11 complete turns (residues 24–150/271–360/456–587/617– 673), surrounded by two small β-barrel domains (β-barrel 1, residues 151–270; β-barrel 2, residues 361–455). Superimposition with the closest homolog, namely PdGH110B (PDB ID: 7JWF; identity: 27.9%)^11)^ aligned 514 Cα atoms out of 810 (RMSD = 1.2 Å), revealing a considerable structural difference (Figure S5). A most striking structural difference of AgaBb is the absence of an α-helix in the β-barrel 2 subdomain, which plays significant structural roles in the dimerization of PdGH110B and formation of the active site in a neighboring subunit (Figure S5B).^11)^ The uncharacterized region observed in the AgaBb844 crystal structure consists of β-sandwich 1 (residues 673–730) and β-sandwich 2 (residues 731–844) (Figure 2B). β-Sandwich 1 is completed by an extension from the core GH110 β-helix domain (residues 588–616). β-Sandwich 2 closely interacts with the core β-helix and β-barrel 1 domain (Figure 2C). A structural homology search using the DALI server^34)^ was performed with each β-sandwich domain, and the only two hits related to CAZymes activity were found with the β-sandwich 1 domain (Table S5): an uncharacterized domain of a carbohydrate-associated hypothetical protein from *Saccharophagus degradans* 2-40 (Z-score = 8.1)^35)^ and a Bacteroidetes-associated carbohydrate-binding often N-terminal (BACON) domain of a xyloglucanase from *Bacteroides ovatus* (Z-score = 8.1).^36)^

### Analysis of the active site

Extensive co-crystallization and soaking experiments with blood group B trisaccharides using a possible catalytic acid residue mutant (D351N) of various constructs did not reveal plausible electron densities in the active site. Although the electron density in the active site of T7-tag_AgaBb673 occupied the position of the –1 subsite (Figure S4B), the orientation of the possible galactose molecule could not be determined owing to low resolution. Therefore, the blood group B trisaccharide model was placed in the active site of AgaBb (Figure 3). In comparison with the PdGH110B structure,^11)^ D351 was suggested to be the catalytic acid residue, whereas D328 and D352 were candidates for the catalytic base residue. The putative catalytic residues are located near the –1 subsite, and R270, D90, D551, and D355 make polar contacts with the non-reducing-end galactose. Galactose in the +1 subsite is stabilized by hydrogen bonds with D351 and D352, and ionic interactions with R475. Fucose in the plausible +1’ subsite interacts with R270 and E380. W509 forms hydrophobic interactions with both the sugars at –1 and +1’ subsites. All these residues are conserved in PdGH110B, except for E380, which is substituted for glutamine.

**Figure 3.**
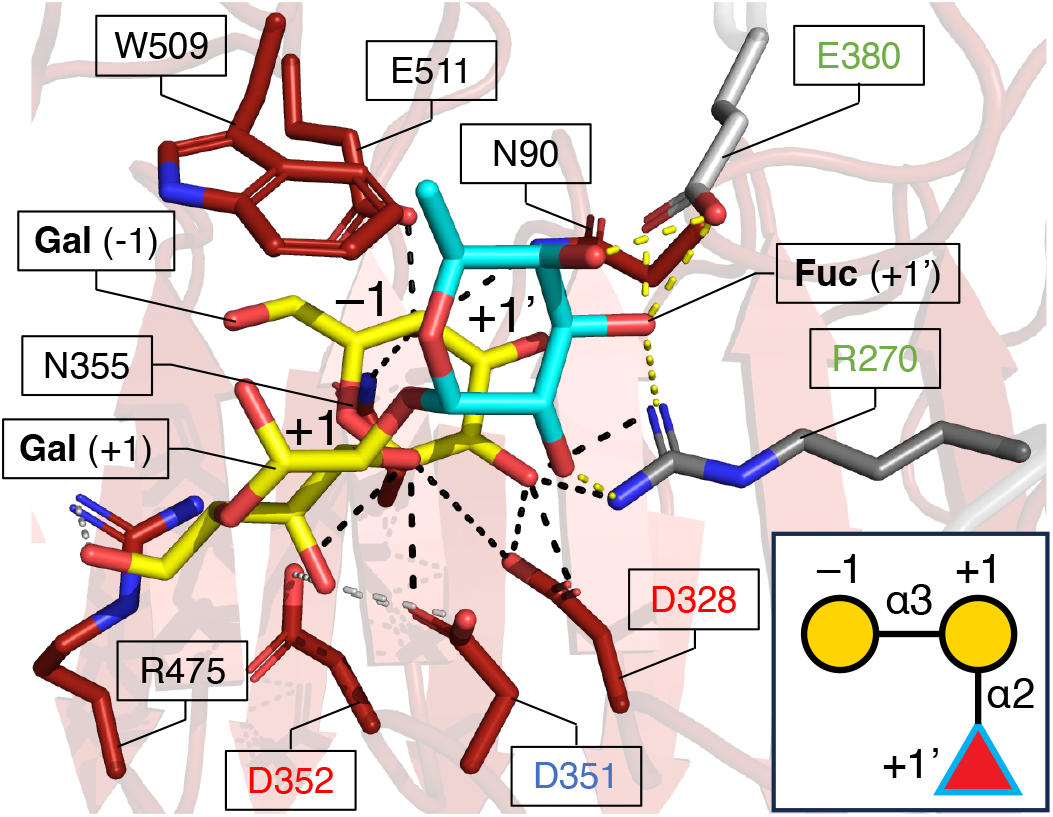
Structure of the active site of AgaBb with modeled blood group B trisaccharide. The protein domains are colored as described in Figure 2. A schematic representation of blood group B trisaccharide is shown in the lower right panel. Gal (yellow) and Fuc (cyan) are shown in sticks and their corresponding subsites in the active site are indicated. The catalytic acid (D351) and catalytic base residue candidates (D328 and D352) are indicated by blue and red characters, respectively. Protein residues constituting the +1’ subsite are indicated by green characters. Polar contacts with the sugar moieties in –1, +1, and +1’ subsites are indicated with black, gray, and yellow dashed lines, respectively.

To validate the potential implication of the active site residues, particularly of fucose recognition at the +1’ subsite, the activity of single substitution mutants toward blood group B trisaccharide was measured using the galactose dehydrogenase-coupling method (Table 1). A mutant of the catalytic acid residue (D351N) exhibited largely decreased activity, similar to that in our previous report.^14)^ The W509A mutant also showed reduced activity, which was comparable to that of D351N, possibly because of the loss of interactions at the –1 and +1’ subsites. R270A mutant in the +1’ subsite retained weak activity. For E380 in the +1’ subsite, an alanine substitution (E380A) was destructive, whereas a conservative substitution with glutamine (E380Q) retained the activity. The results of the mutational analysis support the reliability of the substrate modeling and suggest that correct binding of fucose at the +1’ subsite is important for the activity of AgaBb toward blood group B trisaccharide.

**Table 1.** Relative activity of AgaBb844 mutants toward blood group B trisaccharide.

| **Mutant** | **Subsite** | **Relative Activity (%)<sup>a</sup>** |
| --- | --- | --- |
| D351N | −1 | 1.0 ± 8.8 |
| W509A | −1, +1' | 1.2 ± 3.6 |
| R270A | +1' | 4.7 ± 4.9 |
| E380A | +1' | 4.1 ± 8.9 |
| E380Q | +1' | 30.7 ± 2.7 |

<sup>a</sup> Relative activity compared with the wild-type enzyme (100 ± 1.8%).
Activity toward 2 mM blood group B trisaccharide at pH 6.0 and 37 °C was measured. See Materials and Methods for the detailed assay conditions.

### Thermostability analysis

A thermal shift assay was performed using six C-terminal deletion constructs (Figure 4). In summary, the two shorter constructs (up to residue 730) showed *T*_m_ values <55 °C whereas the four longer constructs (including residues 844 and beyond) exhibited *T*_m_ values >60 °C. The *T*_m_ values of AgaBb673, AgaBb730, AgaBb844, AgaBb945, AgaBb1099, and AgaBb1191 at 20 mg/mL protein concentration were 50.8, 53.4, 61.4, 64.6, 65.0, and 65.7°C, respectively. When the protein concentration was lowered to 15 or 10 mg/mL, a slight increase in *T*_m_ values was observed for every construct. The largest increase in *T*_m_ value (3.9 °C) was observed for AgaBb1191 (difference between 20 mg/mL and 10 mg/mL protein concentration). This result indicates that the β-sandwich 2 domain contributes to the thermostability of AgaBb, and that the oligomeric state is irrelevant.

**Figure 4.**
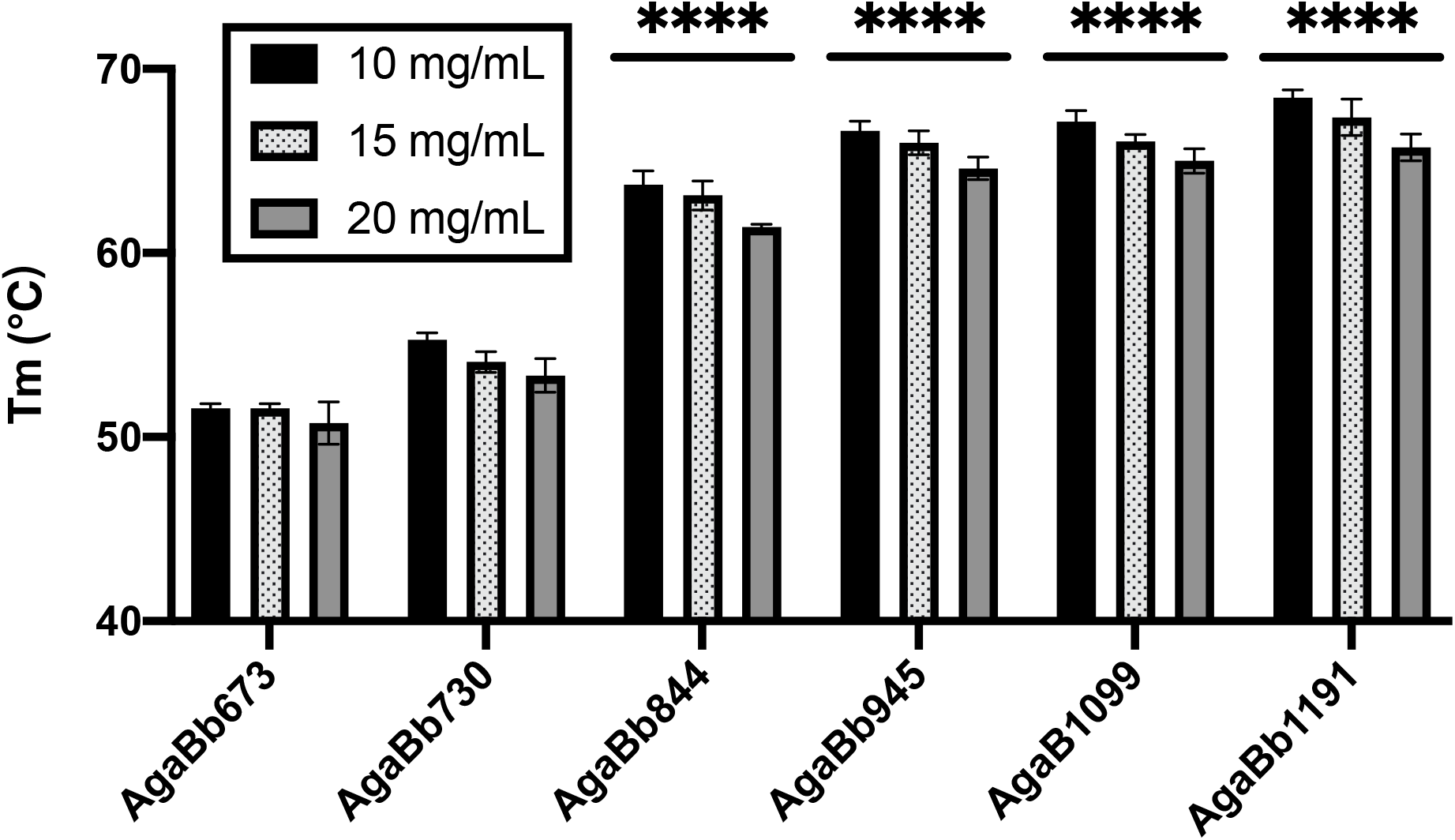
Protein thermostability of C-terminal deletants of AgaBb. Melting temperature (*T*_m_, in °C) of the protein constructs of different lengths at different protein concentrations (10, 15, and 20 mg/mL) were determined by thermal shift assay. Data were collected in triplicate and plotted as the mean ± standard deviation (SD). The average *T*_m_ of the three concentrations of the constructs was compared to that of AgaBb673 using the two-way ANOVA test (****, *P* < 0.0001). Comparison of the average *T*_m_ of the constructs with that of AgaBb730 also displayed similar significance (not shown).

## DISCUSSION

### A mechanism for blood group B antigen recognition

In the present study, we sought to determine the binding mode of a blood group B antigen-specific GH110 α-galactosidase AgaBb and performed mutational analyses to validate the result of substrate modeling. The structural basis of the GH110 catalytic domain and the N-terminal region of the uncharacterized region were clarified. In the active site, we found that R270 and E380 were involved in the recognition of fucose at the +1’ subsite, presumably framing the trisaccharide substrate into the catalytic pocket. Therefore, the fucose residue must be fixed by R270 and E380 prior to anomer-inverting GH catalysis: nucleophilic attack of a water molecule on the –1 subsite galactose facilitated by the catalytic base residue (D352 or D328) and proton donation to the glycosidic bond oxygen by the catalytic acid residue D351. As few residues are involved in forming the +1 and +1’ subsites, the release of the reaction product (H antigen) is expected to be efficient (Figure 3). The conservation of the two key residues involved in fucose recognition (R270 and E380) in the GH110 subfamilies was analyzed by multiple amino acid sequence alignment (Figure 5A). Interestingly, R270 was conserved in all the characterized GH110 enzymes, and E380 is conservatively substituted with glutamine or methionine, with no significant differences between the subfamilies. This observation suggests that other factors may also be involved in the differences in substrate specificity between GH110 subfamilies. To gain further insight, we compared the activesite structures of AgaBb and PdGH110B with the predicted structures of previously characterized GH110 enzymes belonging to subfamilies A and B (Figure 5B).^10,11)^ PdGH110B has more residues around fucose than AgaBb and is more densely packed. The inability of PdGH110B to act on the blood group B antigen may be because of this structural feature at the +1’ subsite, as discussed in a previous report.^11)^ In the predicted active site structures of subfamily A enzymes from *Streptomyces avermitilis, Bacteroides fragilis*, and *Bacteroides thetaiotaomicron* (SaGal110A, BfGal110A, and BtGal110A, respectively), the residues for fucose recognition were almost conserved. In contrast, the predicted active sites of the subfamily B enzymes from *B. fragilis* and *B. thetaiotaomicron* (BfGal110B and BtGal110B, respectively) contained tryptophan, glutamine, and methionine residues that were not found in AgaBb. In addition, the side chain conformations of the amino acids corresponding to R270 differed between subfamilies A and B.

**Figure 5.**
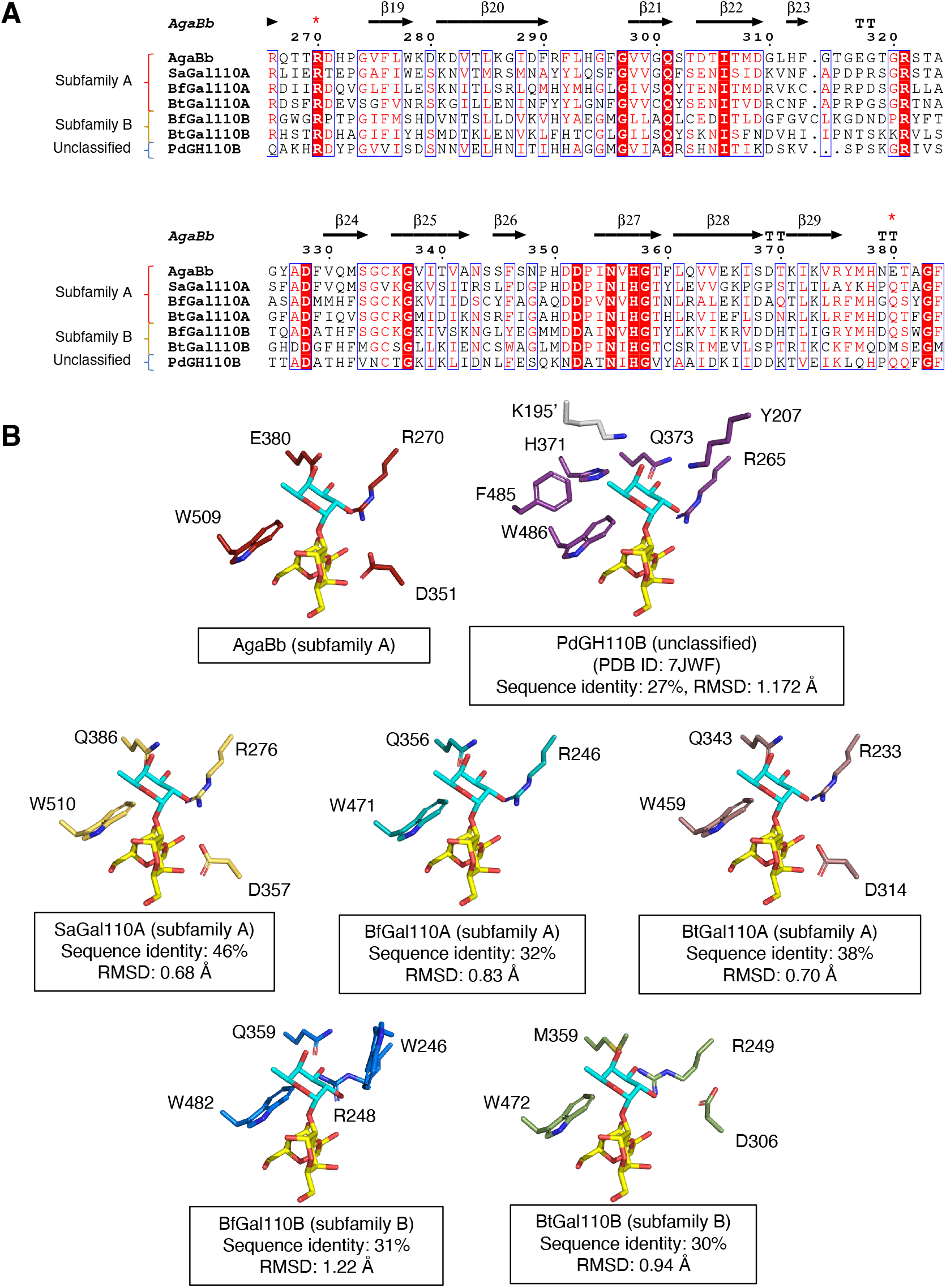
Comparison of the putative fucose recognition site in AgaBb with that of previously characterized GH110 enzymes. (A) Multiple amino-acid sequence alignment. The residues implicated in the fucose recognition in AgaBb are indicated with red asterisks above the sequences. (B) Putative +1’ subsite of AgaBb and GH110 enzymes in subfamilies A and B. Residues within 4 Å distance from the fucose are shown. PdGH110B, α-galactosidase from *P. distincta*; SaGal110A, α-galactosidase from *S. avermitilis*; BfGal110A and BfGal110B, α-galactosidases from *B. fragilis*; BtGal110A and BtGal110B, α-galactosidases from *B. thetaiotaomicron*.

Very recently, the crystal structure of an α-galactosidase from *Akkermansia muciniphila* (AmGH110), which may be active on blood group B antigen and belongs GH110 subfamily A (sequence identity = 27% with AgaBb), was released (PDB ID: 8PVS). Although a related article has not been published yet, we examined the AmGH110 structure. The catalytic domain of AmGH110 also contains β-barrels 1 and 2, and an additional β-sandwich domain is present at the N-terminus (Figure 6A). The sequence identity between AgaBb and AmGH110 is 27.3%, which is slightly lower than the identity between AgaBb and PdGH110B (27.9%). RMSD between AgaBb and AmGH110 was 1.1 Å for 810 Cα atoms, indicating that their backbone structures are similar. In the putative +1’ subsite of AmGH110, R372, Q482, and W596 are conserved (Figure 6B). These structural features may be responsible for the different substrate specificities of the GH110 enzymes.

**Figure 6.**
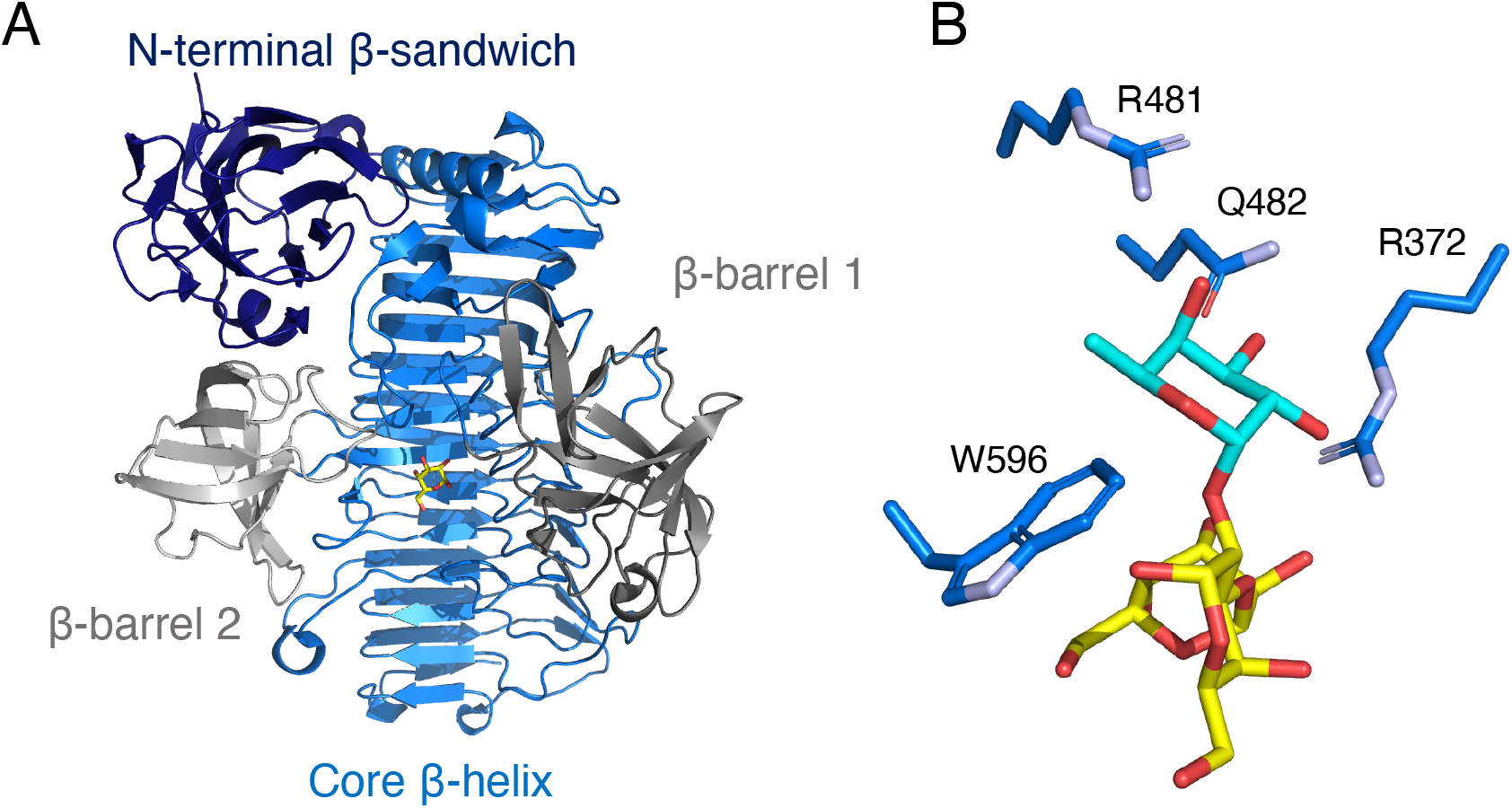
The crystal structure *A. muciniphila* GH110 α-galactosidase (AmGH110) (A) The overall structure. (B) Putative +1’ subsite. Shown as in Figure 6B. The structure available on the database (PDB ID: 8PVS) is shown.

### The uncharacterized region is a protein stabilization factor

In the AgaBb844 structure, the two β-sandwich domains in the uncharacterized region strongly interact with the core β-helix GH110 catalytic domain (Figure 2B). The β-sandwich 2 domain (residues 730–844) forms extensive interactions with the core β-helix and β-barrel 1 domains (Figure 2C). The β-sandwich 1 domain has an integrated fold comprising a part of the uncharacterized region (residues 673–730) and a region extending from the core β-helix (residues 588–616). In the T7-tag_AgaBb700 structure (Figure S4A), the C-terminal part of the core β-helix and an N-terminal part of the β-sandwich 1 domain (residues 618-700) lost most of their secondary structures except for a short β-sheet, and the extended region from the core β-helix (residues 587–617) was disordered. In fact, the thermal shift assay showed a significant increase in the *T*_m_ values for constructs containing the β-sandwich 2 domain (Figure 5), suggesting that the previously uncharacterized region is important for the structural integrity and stabilization of AgaBb.

### Phylogenetic analysis of the uncharacterized region

In the present study, we characterized a region of AgaBb with unknown function. We then attempted phylogenetic analysis to determine the extent to which organisms and proteins conserve this region. A BLAST search using the uncharacterized region of AgaBb (residues 674–945) as a query yielded 102 hits with the lowest similarity scores of sequence identity 31.58%, query cover of 60%, and e-value of 0.026. The phylogenetic tree constructed using this sequence set revealed three major groups (Figure S6). The first group consisted exclusively of *B. bifidum* sequences, of which 44 were phylogenetically very close. The second group was dominated by *Ruminococcus* species of gastrointestinal origin, and contained 22 sequences. The remaining sequences were derived from a wide variety of microorganisms in the animal gut and soil, and their phylogenetic relationships were more ambiguous than those in the first two groups. These sequences were also analyzed using the dbCAN3 meta-server to determine whether they contained CAZyme domains, and 92.3% of the sequences were found to contain the same GH110 + CBM51 domain configuration as that of AgaBb. The CBM2 and CBM54 domains were present in 1.1% of the sequences, whereas 5.5% of the sequences had no functionally annotated domains. Thus, this uncharacterized region may have been acquired independently by *B. bifidum* and *Ruminococcus* sp. to stabilize GH110.

### Feedback from structural prediction as an efficient tool for construct design

The crystal structures presented in this study were determined in the order of T7-tag_AgaBb700, T7-tag_AgaBb673, and then AgaBb844 over a decade. Because crystallographic analysis of AgaBb was challenging, we describe the process briefly. In 2012– 2013, we succeeded in crystallizing the T7-tag_AgaBb700 construct once. We collected several X-ray datasets of the native protein crystals, as well as anomalous diffractions of selenomethionine substituents and heavy atom-soaked crystals. However, we failed to determine the initial phase after extensive SAD trials and multiple isomorphous replacement with anomalous scattering. In 2019, we succeeded in determination of the structure by the SAD method using only anomalous scattering of a platinum derivative dataset, likely because of advances in the phasing pipeline technology of the ccp4 software. The problem of the reproducibility of the crystallization of T7-tag_AgaBb700 prompted us to produce further deletion constructs based on secondary structure prediction using PSIPRED.^37)^ We produced many C-terminal deletions (not shown), but T7-tag_AgaBb673 yielded only low-resolution diffraction-quality crystals. At that time, the homologous structures of the uncharacterized regions had not been reported, and their functions were unknown. In 2021, a highly accurate structure prediction method, AlphaFold2, was released,^38)^ which allowed us to examine the possible domain boundaries of AgaBb. Based on the predicted structural information for the construct design (Figure S1), we succeeded in determining the structure of AgaBb844 six months after the publication of AlphaFold2. We expect that experimental structural biology studies of multidomain enzymes will be further accelerated by state-of-the-art structure prediction tools in the future.

## <List of Abbreviations>

AgaBb: α-galactosidase from *Bifidobacterium bifidum*
Big2: bacterial immunoglobulin-like group 2
BACON: Bacteroidetes-associated carbohydrate-binding often N-terminal
CAZy or CAZymes: carbohydrate-active enzymes
CBM: carbohydrate-binding module
GH: glycoside hydrolase
RMSD: root mean square deviation
SAD: single-wavelength anomalous diffraction
TLC: thin-layer chromatography
*T*_m_: melting temperature

## ACKNOWLEDGMENTS

We thank the staff of the Photon Factory and SPring-8 for the X-ray data collection. We thank Drs. Masahide Hikita and Ayaka Harada for their assistance with this study. We thank Drs Chihaya Yamada, Dohyun Im (Graduate School of Medicine, Kyoto University, Japan), Kazune Tamura (Sir William Dunn School of Pathology, University of Oxford, UK), and Mr. Masanari Tanuma for their experimental support and helpful discussions. This work was supported by JSPS-KAKENHI (24380053 to S.F. and H.A., 23H00322 to S.F.; 8K05494 to H.A.), Mizutani Foundation for Glycoscience, and partially supported by JSPS-KAKENHI (19H00929 and 19K05789 to S.F.) and the Research Support Project for Life Science and Drug Discovery (Basis for Supporting Innovative Drug Discovery and Life Science Research (BINDS)) from AMED under grant number JP22ama121001.

## REFERENCES

1) L. Rydberg: ABO-incompatibility in solid organ transplantation. Transfusion Medicine, vol. 11 (2001).

2) U. Galili: Xenotransplantation and ABO incompatible transplantation: The similarities they share. Transfusion and Apheresis Science, 35, (2006).

3) R. Mitra, N. Mishra and G.P. Rath: Blood groups systems. Indian J Anaesth, 58, 524–528 (2014).

4) Y. Rossez, E. Maes, T. Lefebvre Darroman, P. Gosset, C. Ecobichon, M. Joncquel Chevalier Curt, I.G. Boneca, J.C. Michalski and C. Robbe-Masselot: Almost all human gastric mucin O-glycans harbor blood group A, B or H antigens and are potential binding sites for Helicobacter pylori. Glycobiology, 22, 1193–1206 (2012).

5) G. Misevic: ABO blood group system. Blood and Genomics, 2, 71–84 (2018).

6) P. Rahfeld, L. Sim, H. Moon, I. Constantinescu, C. Morgan-Lang, S.J. Hallam, J.N. Kizhakkedathu and S.G. Withers: An enzymatic pathway in the human gut microbiome that converts A to universal O type blood. Nat Microbiol, 4, (2019).

7) P. Rahfeld and S.G. Withers: Toward universal donor blood: Enzymatic conversion of A and B to O type. Journal of Biological Chemistry, 295, 325–334 (2020).

8) Q.P. Liu, G. Sulzenbacher, H. Yuan, E.P. Bennett, G. Pietz, K. Saunders, J. Spence, E. Nudelman, S.B. Levery, T. White, J.M. Neveu, W.S. Lane, Y. Bourne, M.L. Olsson, B. Henrissat and H. Clausen: Bacterial glycosidases for the production of universal red blood cells. Nature Biotechnology 2007 25:4, 25, 454–464 (2007).

9) E. Drula, M.L. Garron, S. Dogan, V. Lombard, B. Henrissat and N. Terrapon: The carbohydrate-active enzyme database: functions and literature. Nucleic Acids Res, 50, D571–D577 (2022).

10) Q.P. Liu, H. Yuan, E.P. Bennett, S.B. Levery, E. Nudelman, J. Spence, G. Pietz, K. Saunders, T. White, M.L. Olsson, B. Henrissat, G. Sulzenbacher and H. Clausen: Identification of a GH110 subfamily of α1,3-galactosidases: Novel enzymes for removal of the α3Gal xenotransplantation antigen. Journal of Biological Chemistry, 283, (2008).

11) B.E. McGuire, A.G. Hettle, C. Vickers, D.T. King, D.J. Vocadlo and A.B. Boraston: The structure of a family 110 glycoside hydrolase provides insight into the hydrolysis of α-1,3-galactosidic linkages in λ-carrageenan and blood group antigens. Journal of Biological Chemistry, 295, (2020).

12) M. Sakanaka, A. Gotoh, K. Yoshida, T. Odamaki, H. Koguchi, J.Z. Xiao, M. Kitaoka and T. Katayama: Varied pathways of infant gut-associated Bifidobacterium to assimilate human milk oligosaccharides: Prevalence of the gene set and its correlation with bifidobacteria-rich microbiota formation. Nutrients, 12, 71 (2020).

13) T. Katoh, M.N. Ojima, M. Sakanaka, H. Ashida, A. Gotoh and T. Katayama: Enzymatic Adaptation of Bifidobacterium bifidum to Host Glycans, Viewed from Glycoside Hydrolyases and Carbohydrate-Binding Modules. Microorganisms, 8, 481 (2020).

14) T. Wakinaka, M. Kiyohara, S. Kurihara, A. Hirata, T. Chaiwangsri, T. Ohnuma, T. Fukamizo, T. Katayama, H. Ashida and K. Yamamoto: Bifidobacterial α-galactosidase with unique carbohydrate-binding module specifically acts on blood group B antigen. Glycobiology, 23, (2013).

15) M.D. Hansson, K. Rzeznicka, M. Rosenbäck, M. Hansson and N. Sirijovski: PCR-mediated deletion of plasmid DNA. Anal Biochem, 375, (2008).

16) A. D’Arcy, T. Bergfors, S.W. Cowan-Jacob and M. Marsh: Microseed matrix screening for optimization in protein crystallization: What have we learned? Acta Crystallographica Section:F Structural Biology Communications, 70, (2014).

17) W. Kabsch: XDS. Acta Crystallogr D Biol Crystallogr, 66, 125–132 (2010).

18) P.R. Evans and G.N. Murshudov: How good are my data and what is the resolution? Acta Crystallogr D Biol Crystallogr, 69, (2013).

19) P. Skubak, D. Arac, M.W. Bowler, A.R. Correia, A. Hoelz, S. Larsen, G.A. Leonard, A.A. McCarthy, S. McSweeney, C. Mueller-Dieckmann, H. Otten, G. Salzman and N.S. Pannua: A new MR-SAD algorithm for the automatic building of protein models from low-resolution X-ray data and a poor starting model. IUCrJ, 5, (2018).

20) A.J. McCoy, R.W. Grosse-Kunstleve, P.D. Adams, M.D. Winn, L.C. Storoni and R.J. Read: Phaser crystallographic software. J Appl Crystallogr, 40, (2007).

21) K. Cowtan: The Buccaneer software for automated model building. 1. Tracing protein chains. Acta Crystallogr D Biol Crystallogr, 62, (2006).

22) P.D. Adams, P. V. Afonine, G. Bunkóczi, V.B. Chen, I.W. Davis, N. Echols, J.J. Headd, L.W. Hung, G.J. Kapral, R.W. Grosse-Kunstleve, A.J. McCoy, N.W. Moriarty, R. Oeffner, R.J. Read, D.C. Richardson, J.S. Richardson, T.C. Terwilliger and P.H. Zwart: PHENIX: A comprehensive Python-based system for macromolecular structure solution. Acta Crystallogr D Biol Crystallogr, 66, 213–221 (2010).

23) D. Liebschner, P. V. Afonine, M.L. Baker, G. Bunkoczi, V.B. Chen, T.I. Croll, B. Hintze, L.W. Hung, S. Jain, A.J. McCoy, N.W. Moriarty, R.D. Oeffner, B.K. Poon, M.G. Prisant, R.J. Read, J.S. Richardson, D.C. Richardson, M.D. Sammito, O. V. Sobolev, D.H. Stockwell, T.C. Terwilliger, A.G. Urzhumtsev, L.L. Videau, C.J. Williams and P.D. Adams: Macromolecular structure determination using X-rays, neutrons and electrons: Recent developments in Phenix. Acta Crystallogr D Struct Biol, 75, (2019).

24) P. Emsley, B. Lohkamp, W.G. Scott and K. Cowtan: Features and development of Coot. Acta Crystallogr D Biol Crystallogr, 66, (2010).

25) E. Merck: Dyeing Reagents for Thin-Layer and Paper Chromatography. Amines Amino acids Amino acids, (1980).

26) K. Anderson, S.C. Li and Y.T. Li: Diphenylamine-aniline-phosphoric acid reagent, a versatile spray reagent for revealing glycoconjugates on thin-layer chromatography plates. Anal Biochem, 287, (2000).

27) M. Miwa, T. Horimoto, M. Kiyohara, T. Katayama, M. Kitaoka, H. Ashida and K. Yamamoto: Cooperation of β-galactosidase and β-N-acetylhexosaminidase from bifidobacteria in assimilation of human milk oligosaccharides with type 2 structure. Glycobiology, 20, (2010).

28) C. Notredame, D.G. Higgins and J. Heringa: T-coffee: A novel method for fast and accurate multiple sequence alignment. J Mol Biol, 302, (2000).

29) X. Robert and P. Gouet: Deciphering key features in protein structures with the new ENDscript server. Nucleic Acids Res, 42, (2014).

30) J. Zheng, Q. Ge, Y. Yan, X. Zhang, L. Huang and Y. Yin: DbCAN3: automated carbohydrate-active enzyme and substrate annotation. Nucleic Acids Res, 51, W115– W121 (2023).

31) S. Kumar, G. Stecher, M. Li, C. Knyaz and K. Tamura: MEGA X: Molecular evolutionary genetics analysis across computing platforms. Mol Biol Evol, 35, (2018).

32) I. Letunic and P. Bork: Interactive Tree Of Life (iTOL) v4: recent updates and new developments | Nucleic Acids Research | Oxford Academic. Nucleic Acids Research, vol. 47 (2019).

33) E. Krissinel and K. Henrick: Inference of Macromolecular Assemblies from Crystalline State. J Mol Biol, 372, 774–797 (2007).

34) L. Holm: DALI and the persistence of protein shape. Protein Science, 29, (2020).

35) J.H. Hehemann, C. Marsters and A.B. Boraston: Ab initio phasing of a nucleoside hydrolase-related hypothetical protein from Saccharophagus degradans that is associated with carbohydrate metabolism. Proteins: Structure, Function and Bioinformatics, 79, (2011).

36) J. Larsbrink, T.E. Rogers, G.R. Hemsworth, L.S. McKee, A.S. Tauzin, O. Spadiut, S. Klinter, N.A. Pudlo, K. Urs, N.M. Koropatkin, A.L. Creagh, C.A. Haynes, A.G. Kelly, S.N. Cederholm, G.J. Davies, E.C. Martens and H. Brumer: A discrete genetic locus confers xyloglucan metabolism in select human gut Bacteroidetes. Nature, 506, (2014).

37) D.W.A. Buchan and D.T. Jones: The PSIPRED Protein Analysis Workbench: 20 years on. Nucleic Acids Res, 47, (2019).

38) J. Jumper, R. Evans, A. Pritzel, T. Green, M. Figurnov, O. Ronneberger, K. Tunyasuvunakool, R. Bates, A. Žídek, A. Potapenko, A. Bridgland, C. Meyer, S.A.A. Kohl, A.J. Ballard, A. Cowie, B. Romera-Paredes, S. Nikolov, R. Jain, J. Adler, T. Back, S. Petersen, D. Reiman, E. Clancy, M. Zielinski, M. Steinegger, M. Pacholska, T. Berghammer, S. Bodenstein, D. Silver, O. Vinyals, A.W. Senior, K. Kavukcuoglu, P. Kohli and D. Hassabis: Highly accurate protein structure prediction with AlphaFold. Nature, doi:10.1038/s41586-021-03819-2 (2021).

39) P. Rahfeld, L. Sim, H. Moon, I. Constantinescu, C. Morgan-Lang, S.J. Hallam, J.N. Kizhakkedathu and S.G. Withers: An enzymatic pathway in the human gut microbiome that converts A to universal O type blood. Nature Microbiology 2019 4:9, 4, 1475–1485 (2019).

