## Supplementary File for "Crystal structure of *Bifidobacterium bifidum* glycoside hydrolase family 110 α-galactosidase specific for blood group B antigen"

### Supplemental information for Kashima et al.

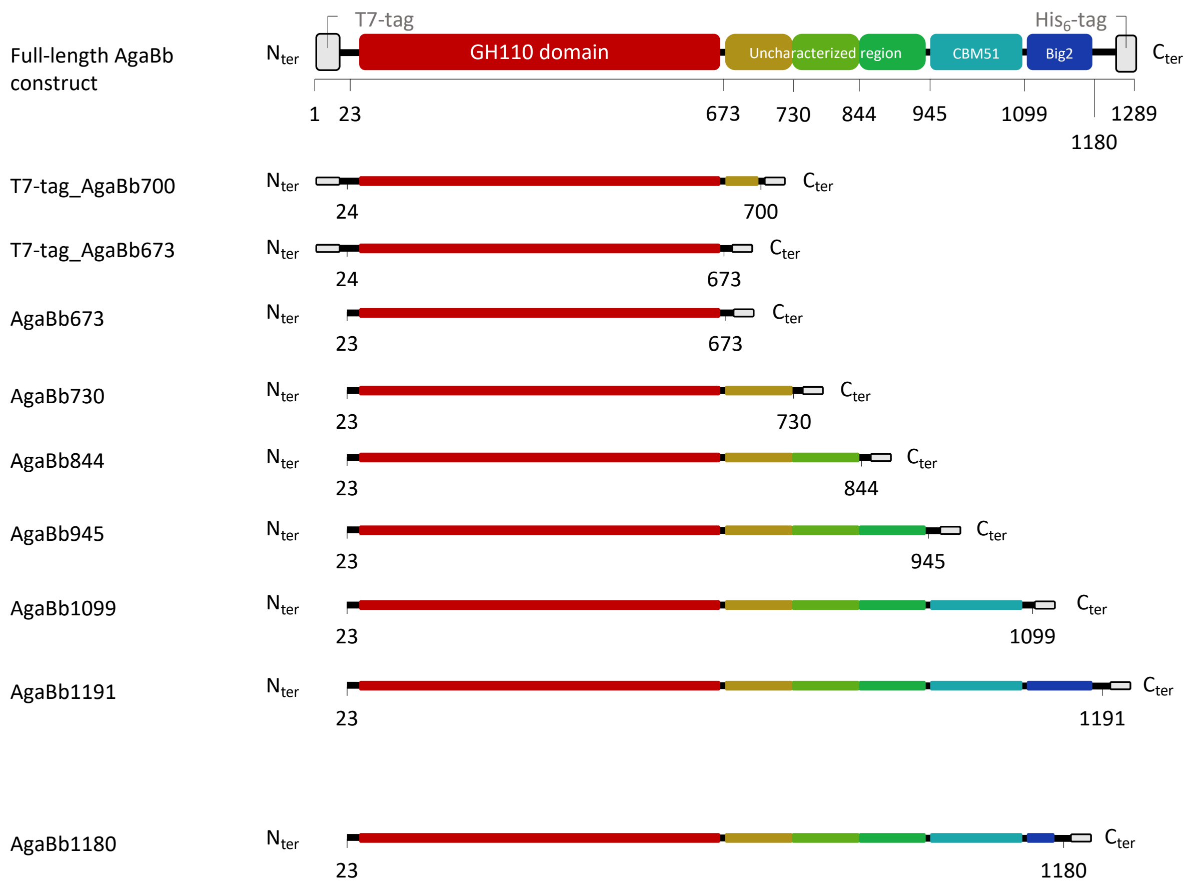


**Figure S1. AgaBb constructs designed and used in the present study.** GH110, glycoside hydrolase family 110; CBM51, carbohydrate-binding module family 51; Big2, bacterial Ig-like domain.

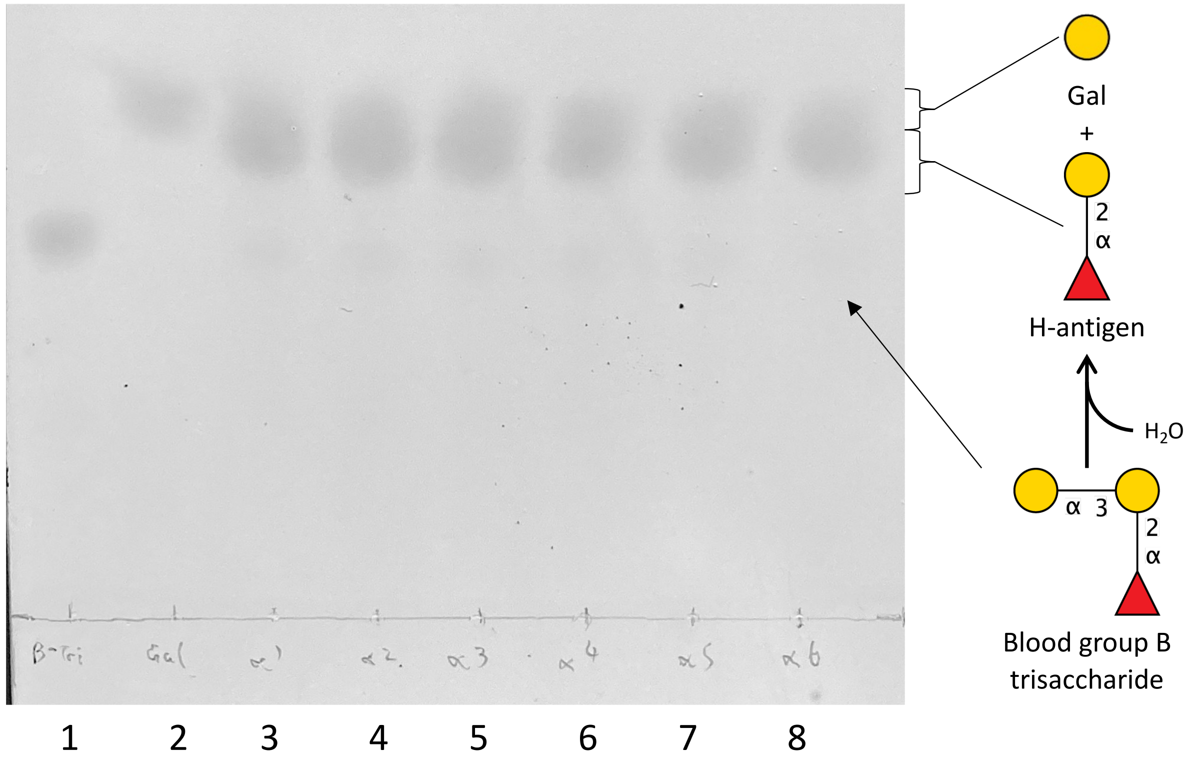


**Figure S2. Activity of the various AgaBb deletion constructs of different lengths toward blood group B trisaccharide.** The substrate (1 mM) was incubated with the purified protein (1 µg/mL) in 50 mM Na-acetate (pH 6.0) overnight at 37 °C. Lanes 1 and 2 correspond to blood group B trisaccharide and galactose used as negative and positive controls, respectively. The enzyme constructs used are AgaBb673 (lane 3), AgaBb730 (lane 4), AgaBb844 (lane 5), AgaBb945 (lane 6), AgaBb1099 (lane 7), and AgaBb1191 (lane 8).

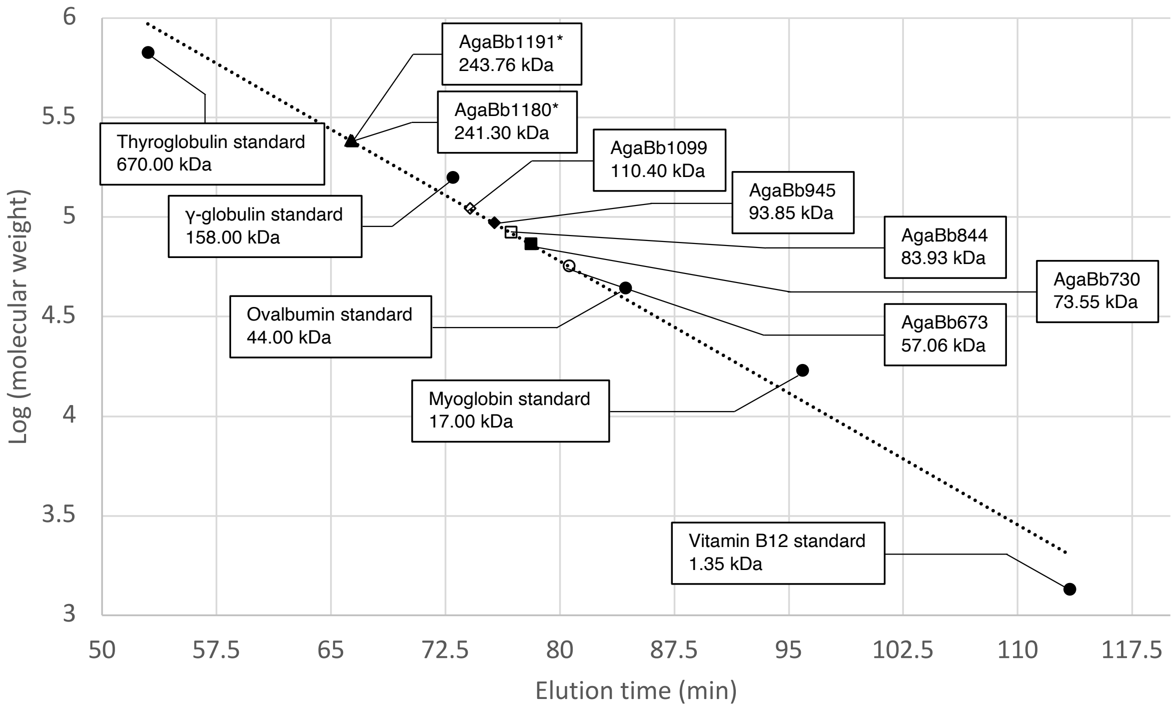


**Figure S3. Gel-filtration chromatography analysis**. The molecular weight of AgaBb constructs was determined by gel-filtration chromatography. The logarithm of the molecular weight is plotted as a function of the elution time. The dots corresponding to the gel filtration standards are shown as closed circles. The calibration line ($R^2$ = 0.978) is shown as a dashed line. Place holders corresponding to AgaBb673 (open circle), AgaBb730 (closed square), AgaBb844 (open square), AgaBb945 (closed rhombus), AgaBb1099 (open rhombus), AgaBb1180 (closed triangle) and AgaBb1191 (open triangle) are shown. AgaBb1191 and AgaBb1180 were measured as dimers and indicated by an asterisk.

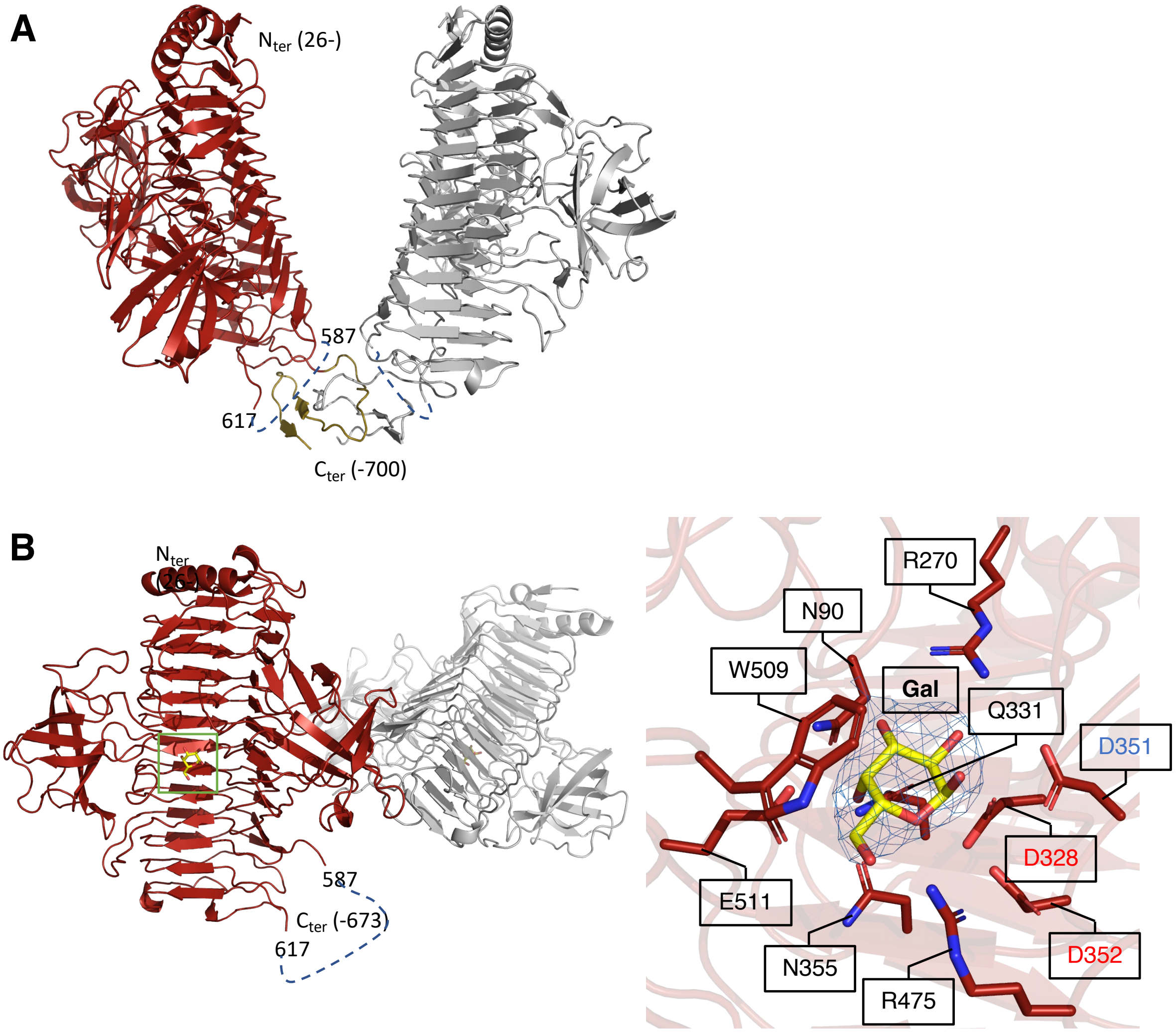


**Figure S4. Structure of T7-tag_AgaBb700 and T7-tag_AgaBb673.** (A) Dimer structure of T7-tag_AgaBb700 in the crystal packing. Residues 587–617 (blue dashed line) were not observed. The C-terminal region (residues 618–700), which corresponds to an N-terminal part of the uncharacterized region of AgaBb, is colored in gold. (B) Two monomers of T7-tag_AgaBb673 in the crystal packing (left) and their active site (right). In the left panel, the active-site area is indicated in the green box. In the right panel, the polder map of Gal (5.0 σ) is shown as a blue mesh. The catalytic acid residue (D351) and catalytic base residue candidates (D328 and D352) are indicated in blue and red characters, respectively.

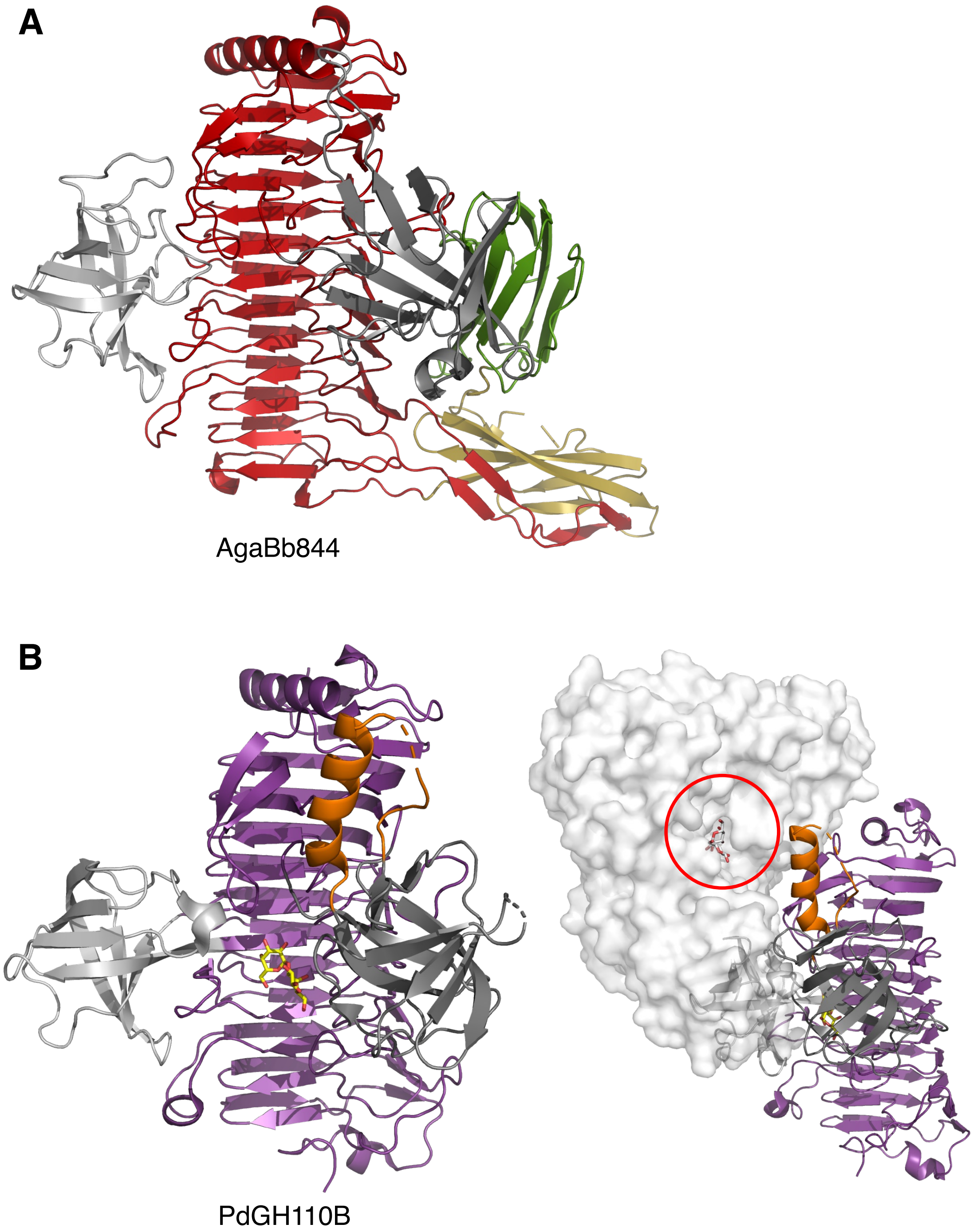


**Figure S5. Structural comparison of AgaBb844 with PdGH110B.** (A) Overall structure of AgaBb844. (B) Monomeric (left) and dimeric (right) structures of PdGH110B (PDB ID: 7JWF). The sequence identity between the catalytic domains of AgaBb and PdGH110B is 27%. Both structures are colored as in Figure 2, except that the core β-helix of PdGH110B is colored in purple. The α-helix present in PdGH110B, but not in AgaBb844, is shown in orange. The galactobiose in the catalytic site of PdGH110B is shown as a yellow stick structure. The dimer counterpart of PdGH110B is shown as a white surface, and the active site is indicated by a red circle (panel B, right)

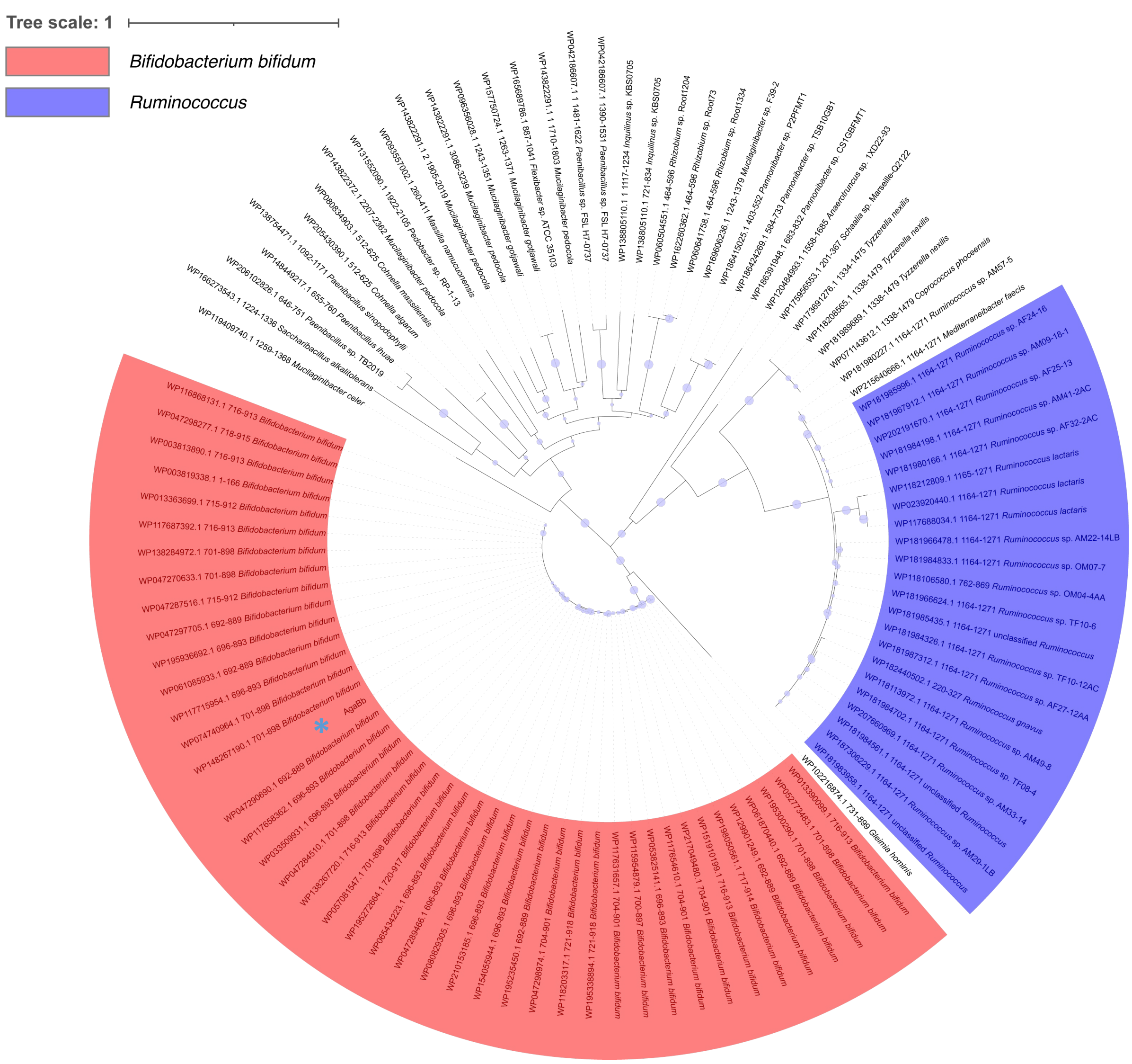


**Figure S6. Phylogenetic tree of the uncharacterized region in AgaBb and its homologs.** The homolog sequences were retrieved by BLAST search using the sequence of residues 674–945 of AgaBb as a query. Bootstrap robustness ranging from 50 to 100% is shown as blue circles of different sizes. AgaBb is indicated with a blue asterisk.

**Table S1. Characteristics and design process of various AgaBb constructs.**

| Construct | Thermal stability [a] | Oligomerization | Crystallizability | Notes |
|---|---|---|---|---|
| T7-tag_AgaBb700 | – | 1 in solution, 2 in the crystal structure | Yes, but hardly reproducible | Used for initial crystallization and X-ray crystallography |
| T7-tag_AgaBb673 | – | 1 | Yes, but poor diffraction quality | Designed following the determination of the T7-tag_AgaBb700 structure |
| AgaBb673 | Low | 1 | Yes, but poor diffraction quality | Designed based on PSIPRED secondary structure prediction[1)], suites with the domain boundary observed on AlphaFold2 structure prediction [2)] |
| AgaBb730 | Low | 1 | No | Designed based on AlphaFold2 structure prediction |
| AgaBb844 | High | 1 | Yes | Designed based on AlphaFold2 structure prediction |
| AgaBb945 | High | 1 | Yes, but poor quality | Designed based on AlphaFold2 structure prediction |
| AgaBb1099 | High | 1 | Yes, but no diffraction | Designed based on PSIPRED secondary structure prediction, suites with the domain boundary observed on AlphaFold2 structure prediction |
| AgaBb1180 | High | 2 | No | Designed based on PSIPRED secondary structure prediction |
| AgaBb1191 | High | 2 | No | Designed based on AlphaFold2 structure prediction |

[a] $T_m$ values below and above 60°C are indicated as low and high, respectively. Thermal stability of constructs indicated by dashes (–) were not measured.

**Table S2. Primer sets used for construct design and site-directed mutagenesis.**

| Objective | Sequence |
|---|---|
| T7-tag deletion | 5’-GAAGGAGATATACATATGGGAGGAGACGTCGTCTCGG-3’<br>5’-CGAGACGACGTCTCCTCCCATATGTATATCTCCTTCTTAAAGTTAAACAAAAT-3’ |
| Δ673 deletion | 5’-GGACGGTCTCGTCCCCCTCGAGCACCACCACCACCA-3’<br>5’-GGTGGTGGTGCTCGAGGGGGACGAGACCGTCCGCGC-3’ |
| Δ730 deletion | 5’-CGGGTGAGCGCCTCGACCCTCGAGCACCACCACCACCACCACTGAGATCCGGCT-3’<br>5’-GTGGTGGTGGTGCTCGAGGGTCGAGGCGCTCACCCGTACGT-3’ |
| Δ844 deletion | 5’-GTGATCGACCGCATTGGCCTCGAGCACCACCACCACCACCACT-3’<br>5’-GTGGTGGTGGTGCTCGAGGCCAATGCGGTCGATCACGAAACGGTAGGT-3’ |
| Δ945 deletion | 5’-GCCGTGACCGGCACGGACCTCGAGCACCACCACCACCACCACTGAGATCCGGCT-3’<br>5’-GTGGTGGTGGTGCTCGAGGTCCGTGCCGGTCACGGCAACC-3’ |
| Δ1099 deletion | 5’-CGGCGTTGAGATGCCGCTCGAGCACCACCACCACCACCACTGAGATCC-3’<br>5’-GGTGGTGGTGCTCGAGCGGCATCTCAACGCCGACGAACCG-3’ |
| Δ1180 deletion | 5’-CGCGGTGCCGTCGGTGCTCGAGCACCACCACCACCACCACTGAGATCCGGCTGC-3’<br>5’-GGTGGTGGTGCTCGAGCACCGACGGCACCGCGACG-3’ |
| Δ1191 deletion | 5’-GTCACGGTGGCGGAGAAACTCGAGCACCACCACCACCACCACT-3’<br>5’-GTGGTGGTGGTGCTCGAGTTTCTCCGCCACCGTGACCGTCACCGAG-3’ |
| D351N mutation | 5’-AGCTTCTCCAACCCGCATAACGACCCGATC-3’<br>5’-GATCGGGTCGTTATGCGGGTTGGAGAAGCT-3’ |

Nucleotide bases involved in the amino-acid substitutions are underlined.

**Table S3. Crystallization conditions.**

| Construct | Protein Concentration (mg/mL) | Reservoir solution | Seed crystal condition | Cryoprotection |
|---|---|---|---|---|
| T7-tag_AgaBb700 | 16.1 | 1.6 M ammonium sulfate, 0.1 M HEPES pH 7.5, 0.1 M NaCl | None | 10% glycerol |
| T7-tag_AgaBb673 | 5 mg/mL | 15% (w/v) PEG 20000, 0.1 M MES-NaOH pH 6.5 | 10% (w/v) PEG 6000, 0.1 M citric acid pH 5.0 (temperature 4 °C) | 5-20% ethylene glycol |
| AgaBb844 | 37.5 mg/mL | 10% (w/v) PEG 3350, 0.1 M sodium iodide | 10% (w/v) PEG 6000, 0.1 M citric acid pH 5.0 (protein:reservoir ratio: 1:2) | 5-20% PEG 300 |

**Table S4. Crystallographic data collection and refinement statistics of AgaBb.**

| Data set | T7-tag_AgaBb700 + $K_2PtCl_4$ | AgaBb844 | T7-tag_AgaBb700 | T7-tag_AgaBb673 |
|---|---|---|---|---|
| Data collection [a] | | | | |
| Beamline | KEK PF NW12A | KEK PF BL1A | KEK-PF NW12A | KEK PF BL1A |
| Wavelength (Å) | 1.0715 | 1.0520 | 1.0000 | 1.0800 |
| Space group | $P2_12_12_1$ | $C2$ | $P2_12_12_1$ | $P2_1$ |
| Unit cell | | | | |
| a, b, c (Å) | 71.70, 126.58, 190.95 | 127.69, 68.77, 291.31 | 70.56, 125.06, 196.79 | 63.34, 73.15, 177.97 |
| β (°) | 90.00 | 91.40 | 90.00 | 91.38 |
| Resolution (Å) | 47.74–3.15 (3.32–3.15) | 48.59–2.02 (2.06–2.02) | 48.04–1.96 (1.99–1.96) | 47.88-3.50 (3,83-3.50) |
| Total reflections | 451,817 (66,099) | 1,146,655 (56,454) | 941,393 (46,166) | 71,714 (16,916) |
| Unique reflections | 30.804 (4,425) | 161,638 (7,796) | 126,369 (6,186) | 20,905 (4,934) |
| $CC_{1/2}$ | 1.00 (0.92) | 1.00 (0.88) | 1.00 (0.90) | 0.76 (0.35) |
| Completeness (%) | 100(100) | 98.2 (96.9) | 100 (100) | 99.9 (100) |
| Multiplicity | 14.7 (14.9) | 7.1 (7.2) | 7.4 (7.5) | 3.4 (3.4) |
| Mean $I/\sigma(I)$ | 18.8 (3.8) | 8.9 (2.7) | 15.1 (3.0) | 2.9 (1.7) |
| $R_{merge}$ | 0.138 (0.899) | 0.124 (0618) | 0.088 (0.643) | 0.377 (0.813) |
| Mol/ASU [b] | | 3 | 2 | 2 |
| Refinement | | | | |
| Resolution (Å) | | 48.63–2.02 | 46.84–1.96 | 47.92-3.50 |
| No. of reflections | | 161,637 | 126,272 | 20,893 |
| $R_{work}/R_{free}$ [c] | | 0.190/0.229 | 0.167/0.202 | 0.211/0.296 |
| Number of atoms | | | | |
| Amino acids | | 18,216 | 9,827 | 9,431 |
| Ions | | 5 | 1 | - |
| Ligands | | - | 23 | 48 |
| Waters | | 792 | 1193 | - |
| B-factors (Å$^2$) | | | | |
| Amino acids | | 34.46 | 33.36 | 24.93 |
| Ions | | 12.01 | 50.92 | - |
| Ligands | | - | 56.90 | 34.57 |
| Waters | | 34.45 | 44.56 | - |
| RMSD from ideal values | | | | |
| Bond lengths (Å) | | 0.0070 | 0.0084 | 0.0045 |
| Bond angles (°) | | 1.344 | 1.463 | 1.091 |
| Ramachandran plot (%) | | | | |
| Favored | | 94.6 | 95.0 | 87.3 |
| Allowed | | 4.2 | 3.6 | 9.3 |
| Outlier | | 1.2 | 1.4 | 3.3 |
| PDB code | | 8YK1 | 8YK2 | 8YK3 |

[a] Values in parentheses are for the highest-resolution shell.

[b] Number of molecules per asymmetric unit

[c] $R_{free}$ was calculated for a randomly chosen 5% of the reflections that were not used for structure refinement, and $R_{work}$ was calculated for the remaining reflections.

**Table S5. Results of the structural similarity search using the DALI server**

| Protein | Source organism | PDB (chain) | Z score | RMSD (Å) | $N_{align}$[a] | $\%_{seq}$[b] |
|---|---|---|---|---|---|---|
| β-sandwich 1 (588-616/673-730) | | | | | | |
| Surface layer protein | *Geobacillus stearothermophilus* | 4IUD (A) | 10.5 | 1.7 | 78 | 27 |
| Tail tube protein | *Escherichia* virus T5 | 5NGJ (A) | 9.6 | 2.7 | 84 | 14 |
| Intimin | *Escherichia coli* O127:H6 | 6TQD (B) | 9.2 | 2.4 | 84 | 8 |
| Major tail protein V | *Escherichia* virus Lambda | 2L04 (A) | 8.8 | 2.2 | 78 | 21 |
| Collagenase | *Hathewaya histolytica* | 2Y72 (B) | 8.6 | 2.2 | 79 | 10 |
| Cadherin-5 | *Mus musculus* | 3LND (C) | 8.4 | 2.5 | 82 | 12 |
| Protocadherin-15 | *Homo sapiens* | 4XHZ (A) | 8.3 | 2.5 | 83 | 14 |
| Cadherin-11, cadherin-6 chimera | *Mus musculus* | 6CGB (A) | 8.2 | 2.6 | 82 | 12 |
| Cadherin-6 | *Mus musculus* | 3LND (D) | 8.2 | 2.6 | 83 | 12 |
| Cadherin-7 | *Mus musculus* | 6CGS (A) | 8.2 | 2.5 | 82 | 15 |
| Vascular endothelial cadherin | *Gas gallus* | 3PPE (B) | 8.2 | 2.7 | 82 | 16 |
| S-layer protein SAP | *Bacillus anthracis* | 6HHU (A) | 8.2 | 2.5 | 83 | 20 |
| Carbohydrate-associated hypothetical protein | *Saccharophagus degradans* 2-40 | 2YHG (A) | 8.1 | 2.3 | 79 | 10 |
| Xyloglucanase | *Bacteroides ovatus* | 3ZMR (B) | 8.1 | 2.4 | 74 | 14 |
| Cadherin-11 | *Mus musculus* | 2A4E (A) | 8.1 | 2.7 | 82 | 13 |
| Cadherin-10 | *Mus musculus* | 6CG6 (A) | 8.1 | 2.5 | 82 | 13 |
| Dyslexia-associated protein KIAA0319-like protein | *Mus musculus* | 6NZ0 (Z) | 8.1 | 2.7 | 79 | 10 |
| Nidogen-1 | *Mus musculus* | 1GL4 (B) | 8.0 | 2.2 | 76 | 12 |
| β-sandwich 2 (731-844) | | | | | | |
| Chromatin structure-remodeling complex subunit RSC4 | *Saccharomyces cerevisiae* S288C | 6V8O (H) | 10.9 | 1.9 | 91 | 11 |

[a] Number of aligned residues

[b] Sequence identity
